# Purinergic signaling is enhanced in the absence of UT-A1 and UT-A3

**DOI:** 10.1101/663252

**Authors:** Nathaniel J. Himmel, Richard T. Rogers, Sara K. Redd, Yirong Wang, Mitsi A. Blount

## Abstract

ATP is an important paracrine regulator of renal tubular water and urea transport. The activity of P2Y_2_, the predominant P2Y receptor of the medullary collecting duct, is mediated by ATP, and modulates urinary concentration. To investigate the role of purinergic signaling in the absence of urea transport in the collecting duct, we housed wild-type (WT) and UT-A1/A3 null (UT-A1/A3 KO) mice in metabolic cages to monitor urine output, and collected tissue samples for analysis. We confirmed that UT-A1/A3 KO mice are polyuric, and concurrently observed lower levels of urinary cAMP as compared to WT, despite elevated serum vasopressin (AVP) levels. Because P2Y_2_ inhibits AVP-stimulated transport by dampening cAMP synthesis, we suspected that, similar to other models of AVP-resistant polyuria, purinergic signaling is increased in UT-A1/A3 KO mice. In fact, we observed that both urinary ATP and purinergic-mediated prostanoid (PGE_2_) levels were elevated. Tissue analysis shows that P2Y_2_ mRNA levels remain unchanged, while P2Y_2_ protein levels are elevated in KO mice. Collectively, our data suggest that the reduction of medullary osmolality due to the lack of UT-A1 and UT-A3 induces an AVP-resistant polyuria that is possibly exacerbated by, or at least correlated with, enhanced purinergic signaling.

**Summary statement:** Physiological analyses suggest that mice lacking urea transporters have increased renal purinergic signaling, perhaps contributing to previously observed vasopressin-resistant polyurias.

## Introduction

A hypertonic inner medulla is necessary for concentrating urine during antidiuresis, and urea is essential to this mechanism (Sands et al., 2011). Along with sodium and chloride, urea is a major contributor to the increased osmolality of the inner medullary interstitium (Sands and Layton, 2009). High interstitial osmolality is sustained by urea reabsorption in the inner medullary collecting duct (IMCD), which in turn drives water reabsorption through the aquaporin channel AQP2 (Klein et al., 2010; Pannabecker et al., 2008; Sands and Layton, 2009). While urea is freely membrane permeable and, with sufficient time, will equalize across a lipid bilayer, maintenance of a hypertonic inner medulla requires rapid urea reabsorption in the IMCD(Sands, 2003). This is facilitated by the urea transport proteins UT-A1 and UT-A3 (Klein et al., 2011).

Arginine vasopressin (AVP) is the chief regulator of water reabsorption in the collecting duct. Antidiuresis is initiated by AVP binding to the G protein-coupled receptor vasopressin receptor 2 (V2R), which leads to increased intracellular levels of cAMP (Rieg et al., 2010; Vallon and Rieg, 2011). Elevated cAMP levels promote phosphorylation of UT-A1 and AQP2, increasing UT-A1-mediated urea reabsorption, and thereby AQP2-mediated water entry (Blount et al., 2008; Hoffert et al., 2006). In the presence of a transepithelial osmolality gradient, AVP induces IMCD cell swelling (Chou et al., 2008; Ganote et al., 1968), which releases local factors such as the purine nucleotide adenoside-5’-triphosphate (ATP), augmenting the AVP response (Vallon and Rieg, 2011).

Renal epithelial cells release ATP in response to multiple stimuli, including increased luminal flow rate and elevated cell volume (Schwiebert, 2001; Vallon and Rieg, 2011). Once released into the extracellular environment of the inner medulla, ATP acts locally on membrane-bound P2 purinergic receptors. Several receptor subtypes – including both ionotropic P2X receptors and metabotropic P2Y receptors – are localized to the rat collecting duct (Wildman et al., 2009). In humans, of the seven known P2X receptor subunits, only P2X_4_ has been detected in significant amounts in the collecting duct (Chabardes-Garonne et al., 2003). Studies have shown that broad activation of P2X receptors increases water excretion; however, these effects were largely attributed to the proximal tubule (Jankowski et al., 2011). Although evidence suggests that P2X receptors are important to various aspects of renal physiology, the specific role P2X receptors play in the IMCD has not yet been elucidated (Burnstock et al., 2014; Unwin et al., 2003).

Several studies have demonstrated that extracellular purines modulate water handling in the IMCD through P2Y receptors (Boone and Deen, 2008; Burnstock et al., 2014). These G-protein-coupled receptors are subdivided based on the targets of their associated G-protein subunits; of the eight P2Y receptors identified, P2Y_1_, P2Y_2_, P2Y_4_, P2Y_6_, and P2Y_11_ stimulate phospholipase C through the G-protein Gαq (Erb et al., 2006; Vallon and Rieg, 2011). Although many P2Y receptors have been reported in renal epithelia and cultured cell lines of renal origin, the P2Y_2_ subtype is apparently the most predominant P2Y receptor expressed in native collecting duct tissue (Insel et al., 2001; Kishore et al., 2000; Schwiebert and Kishore, 2001; Turner et al., 2003).

Following the identification of the receptor’s location in the collecting duct, studies revealed that ATP-driven P2Y_2_ receptor activation is an important regulator of water reabsorption (Ecelbarger et al., 1994; Edwards, 2002; Kishore et al., 1995; Rouse et al., 1994; Sun et al., 2005a; Vallon and Rieg, 2011). As discussed above, elevated cAMP levels promote AQP2-mediated water entry, increasing cell volume (Blount et al., 2008; Hoffert et al., 2006). Increases in cell volume lead to the release of ATP at both the basolateral and apical sides of the IMCD (Hovater et al., 2008), and subsequent ATP driven P2Y_2_ receptor activation inhibits AVP-mediated water permeability via phospholipase C inhibition of cAMP, thereby normalizing cell volume and accelerating the excretion of free water (Ecelbarger et al., 1994; Edwards, 2002; Kishore et al., 1995; Rouse et al., 1994; Sun et al., 2005a; Vallon and Rieg, 2011). Experiments in isolated IMCD indicate that ATP can stimulate the release of prostaglandin E_2_ (PGE_2_) by activation of P2Y_2_ receptors (Sun et al., 2005a; Welch et al., 2003). PGE_2_ is known to affect the transport of water and urea in the IMCD (Rouch and Kudo, 2000), presumably through interactions of a purinergic system, by tonically inhibiting the stimulating effect of AVP in the renal medulla.

While not fully understood, evidence does suggest that urea transport in the IMCD is regulated through purinergic signaling. In mice lacking P2Y_2_ receptors, urea transporter expression is elevated, which likely contributes to the observed increased urinary concentration capacity of these animals (Zhang et al., 2008). Furthermore, in animals with elevated inner medullary purinergic signaling, UT-A1 protein abundance is reduced (Blount et al., 2010; Zhang et al., 2009), and functional data demonstrate that urea transport can be modulated by PGE_2_ post-cAMP-dependent events (Rouch and Kudo, 2000). However, how urea transport might affect the purinergic-prostanoid system remains to be fully elucidated. In this study, we aimed to elucidate the importance of urea transport to purinergic signaling in the IMCD using transgenic mice lacking UT-A1 and UT-A3. We show that, in the absence of UT-A1 and UT-A3, there are unexpectedly low levels of luminal cAMP, despite elevated vasopressin levels. Levels of ATP, PGE2, and P2Y_2_ protein expression in the IMCD are increased, collectively suggesting an increase in purinergic receptor activation in the absence of urea transport.

## Materials and Methods

### Animals

All animal protocols were approved by the Emory University Institutional Animal Care and Use Committee. All studies were performed using male mice between 8-10 weeks of age that either (1) did not express UT-A1 or UT-A3 (UT-A1/A3^−/−^; UT-A1/A3 KO) or (2) were control, wild-type littermates (UT-A1/A3^+/+^; WT). UT-A1/A3 KO mice were a generous gift from Dr. Jeff M. Sands (Emory University, Atlanta, GA) who regenerated the line on a C56BL/6J background from the original NIH source, as previously described (Fenton et al., 2004; Ilori et al., 2013).

Mice were individually housed in Tecniplast Single Metabolic Cages (Tecniplast USA Inc, Exton, PA) for 24 hours of acclimation, followed by 24 hours of urine collection and food/fluid intake measurements.

### Analysis of urine and serum samples

Urine osmolality was measured using a Wescor 5520 Vapor Pressure Osmometer (Wescore, Logan, UT). Urine creatinine was measured by the Jaffe reaction with a colorimetric kit (BioVision, Inc., Milpitas, CA). Urinary levels of cAMP, ATP and prostaglandin were measured using a Cyclic AMP EIA kit (Caymen Chemical, Ann Arbor, MI), a Prostaglandin E2 Express EIA kit (Caymen Chemical, Ann Arbor, MI), and a colorimetric ATP assay kit (BioVision, Inc., Milpitas, CA), respectively. Serum vasopressin levels were measured using an Arg^8^-Vasopressin ELISA Kit (Enzo Life Sciences,). All assay kits were used according to manufacturer specifications.

### Western blot analysis

Whole kidneys were collected, dissected into inner medulla (IM) and outer medulla (OM), and lysates prepared using an SDS lysis buffer. Protein content was quantified using the Bio-Rad DC Protein Assay Kit (Bio-Rad Laboratories, Hercules, CA). Proteins (20 µg/lane) were resolved on 12.5% SDS-PAGE gels and then electroblotted to polyvinylidene difluoride (PVDF) membranes (Millipore, Bedford, MA). After blotting, membranes were blocked for 1 hour in Odyssey Blocking Buffer (Li-COR, Lincoln, NE) and then probed overnight at 4°C with a primary antibody specific against UT-A1 (1:1000; previously characterized (Naruse et al., 1997)), UT-A3 (1:100; previously characterized (Blount et al., 2007)), AQP2 (1:2000; StressMarq, Victoria, BC, Canada), or P2Y_2,_ (1:200; Alomone Labs, Jerusalem, Israel). Membranes were washed in TBS-tween solution, then probed with Alexa680-conjugated anti-rabbit secondary antibody (1:4000; Life Technologies, Grand Island, NY). Probed membranes were again washed, and then visualized using infrared detection via the LI-COR Odyssey protein analysis system. To ensure equal loading and to quantify densitometric scanning, membranes were stained with coomassie blue to visualize total protein load. All densitometries were collected using ImageJ software (National Institutes of Health, Bethesda, MD).

### Immunofluroescence

Tissues were prepared as described previously (Sim et al., 2014). Whole kidney sections were hydrated with ethanol, and endogenous peroxides were quenched with 3% H_2_O_2._ After antigen retrieval using TEG buffer and quenching free aldehyde groups with 50 mM NH_4_Cl/PBS, sections were blocked in 1% BSA/PBS for 30 min at room temperature. Sections were incubated overnight at 4°C with an antibody specific to AQP2 (1:300, Santa Cruz Biotechnology, Dallas, Texas). After several PBS washes, sections were incubated for 2 hours at room temperature with Alexa Fluor 456 conjugated donkey anti-goat secondary antibody (1:200, Invitrogen, Carlsbad, CX). After several PBS washes, sections were mounted with ProLong Gold Antifade with DAPI (Cell Signaling, Danvers, MA), and observed under an Olympus IX71 inverted microscope.

### Real Time qRT-PCR

IM tissue was collected and stored in RNAlater solution (Life Technologies, Carlsbad, CA) until homogenization. Purified RNA was extracted using PureLink RNA Mini Kit (Ambion – Life Technologies, Carlsbad, CA), and quantified via UV spectrophotometry. Taqman RT-PCR reagents (Applied Biosystems, Carlsbad, CA) were used to generate all cDNA. Using predeveloped Taqman assays, real-time quantitative PCR was performed for P2Y_2_ and the housekeeping transcript 18S (Applied Biosystems, Carlsbad, CA). PCR was performed with 20ng of cDNA and the corresponding primer pairs using an iCycler Real-Time Detection System (BioRad Laboratories, Hercules, CA). The relative quantity of mRNA was determined using the comparative Ct method and expression levels of P2Y_2_ receptor mRNA were normalized to ribosomal 18S expression.

### Statistical analysis

Data are expressed as the mean ± standard error of the mean, and n is the number of animals. Differences were determined by Student’s t-test using GraphPad Instat Software (La Jolla, CA). P values <0.05 were considered statistically significant.

## Results

### UT-A1/A3 KO mice are polyuric and have a reduced ability to reabsorb urea

Western blot analysis of inner medullary tissues collected from the UT-A1/A3 null strain developed by Ilori et al. confirms that UT-A1/A3 KO mice do not express either UT-A1 or UT-A3 proteins (Fig. 1a) (Ilori et al., 2013). Inner medullary AQP2 protein expression was unaffected by the loss of urea transporters under basal conditions (Fig. 1b), and immunofluorescence tagging confirms that AQP2 was present at the apical membrane of IMCD cells in both WT and UT-A1/A3 KO animals (Fig. 2).

**Fig. 1.**
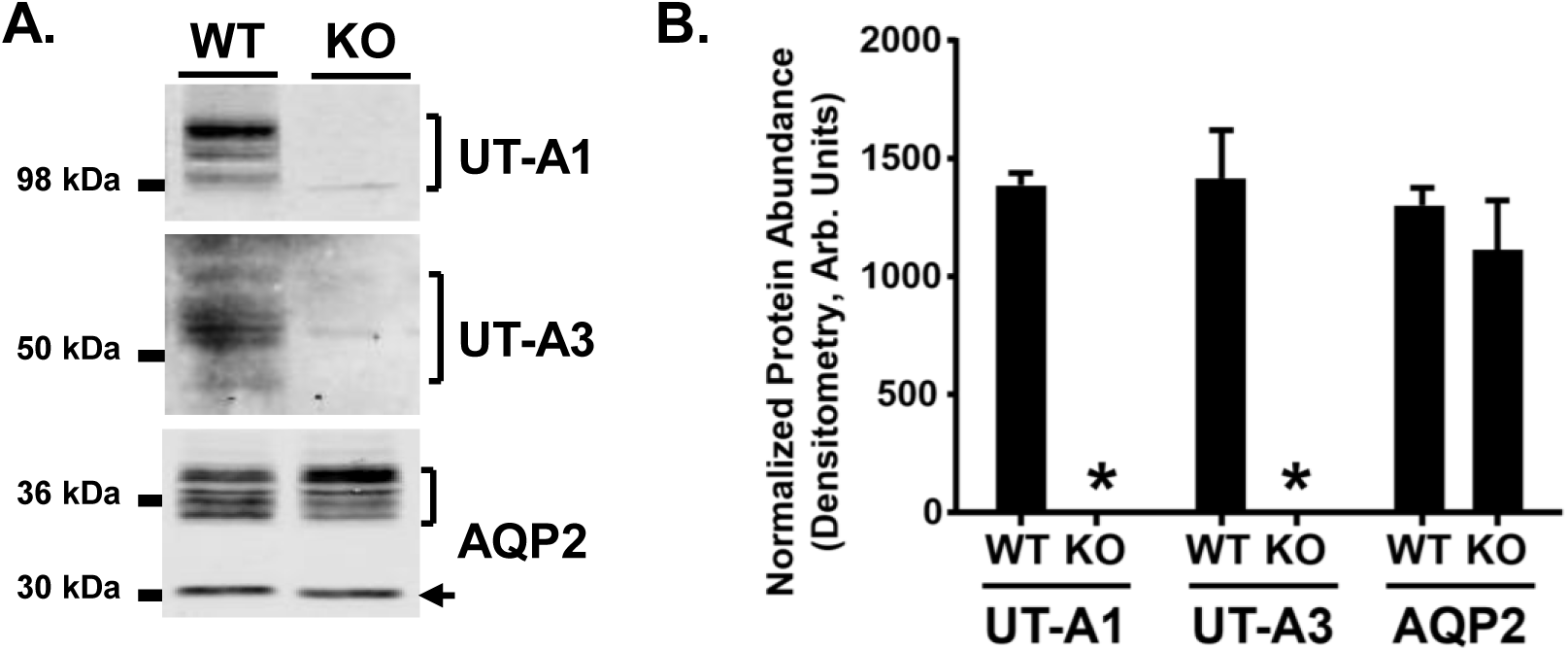
AQP2 is unchanged in the inner medulla of UT-A1/A3 KO mice. Inner medullary tissue was dissected from control, wild-type mice (UT-A1/A3+/+; WT) and UT-A1/UT-A3 null mice (UT-A1/A3-/-; KO). **a** Representative western blot images from tissues probed for UT-A1, UT-A3, and AQP2. Brackets indicate the positive identification of the glycosylated proteins. The arrow indicates the un-glycosylated form of AQP2. **b** Cumulative protein abundance quantification by densitometry presented as means ± SEM. Differences in abundances were determined using a Student’s t-test where p < 0.05 was significant (*); n = 6

**Fig. 2.**
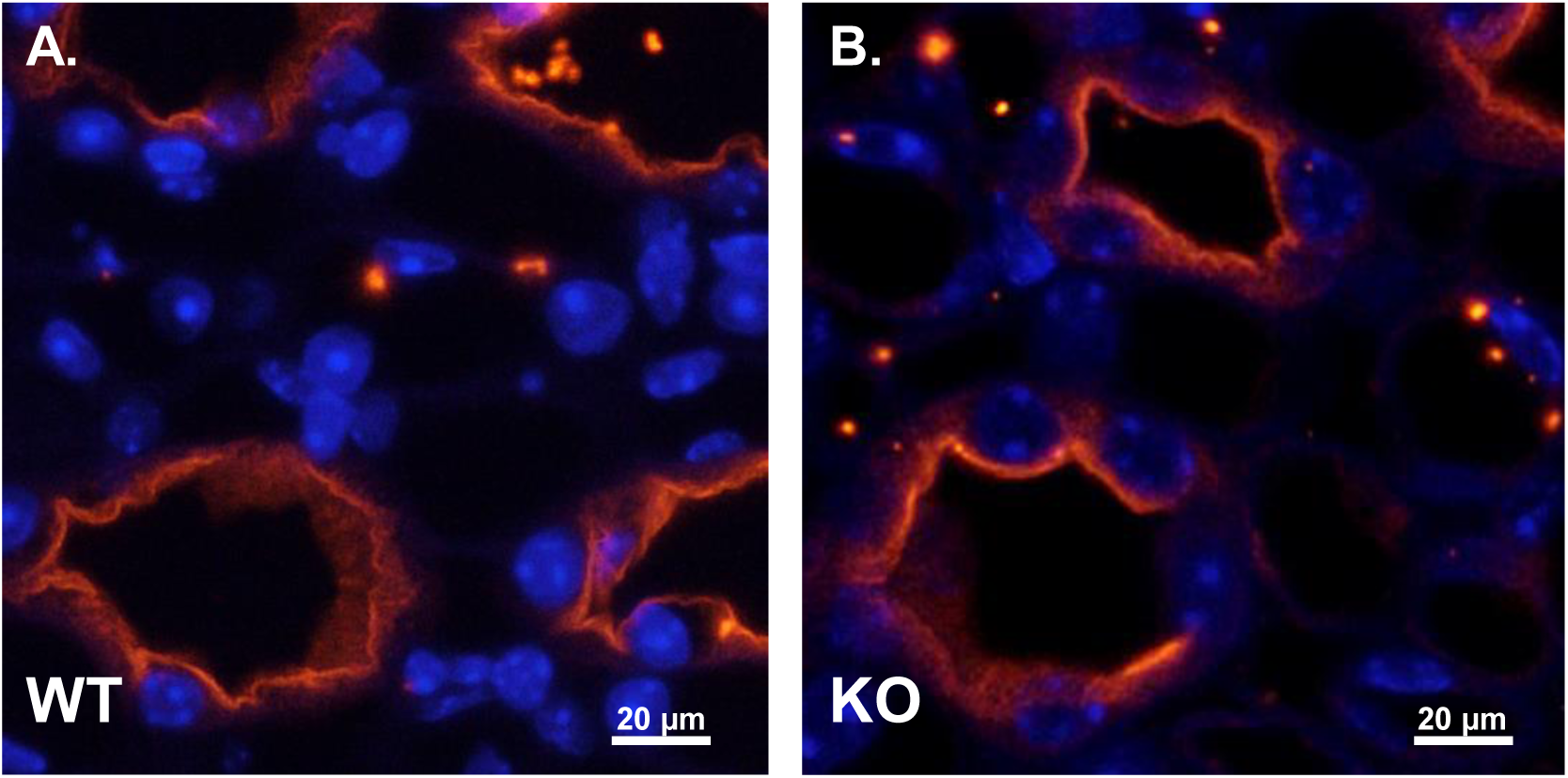
AQP2 localization in the IMCD is similar in WT and UT-A1/A3 KO mice. Perfusion-fixed kidneys were collected from wild-type mice (WT) and UT-A1/A3 KO mice (KO). Following paraffin embedding, 4 µm-thick sections were stained for AQP2 (red) and DAPI (blue). Shown are representative images taken from the inner medulla of WT (**a**) and KO (**b**) mice. n = 3

Compared to WT mice, UT-A1/A3 KO mice had an elevated 24-hour urine volume (Fig. 3a) and a much lower urine osmolality (Fig. 3b), demonstrating that they were polyuric. When corrected for urine dilution, urinary urea concentrations were roughly 7.5-fold higher in UT-A1/A3 KO mice compared to WT animals (Fig. 3c), demonstrating a decreased ability to reabsorb urea in the absence of UT-A1 and UT-A3. This metabolic characterization of UT-A1/A3 KO mice confirms previous reports establishing the importance of UT-1 and UT-A3 in urine concentration (Fenton et al., 2004; Ilori et al., 2013).

**Fig. 3.**
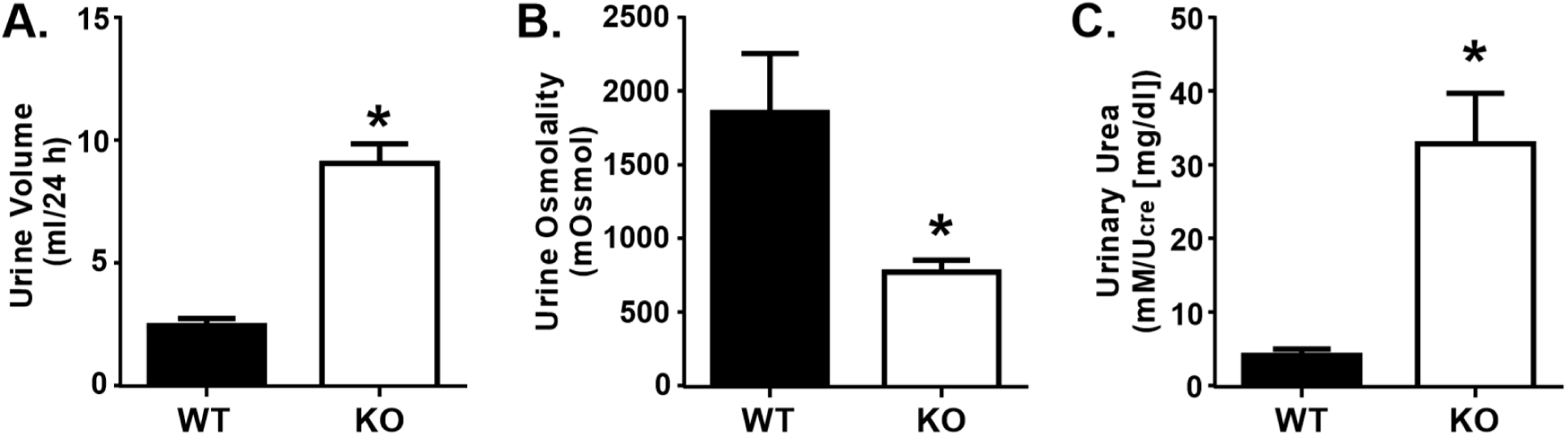
UT-A1/A3 KO mice are polyuric. Control, wild-type mice (WT) and UT-A1/A3 KO mice (KO) were housed in metabolic cages and urine was collected over 24 hours. **a** Total volume of urine collected under oil during the timed 24-hour period. **b** Urine osmolality of the collected urine. **c** Urinary urea measured from collected urine normalized to creatinine (U_cre_) to correct for differences in urine dilution. All data presented as means ± SEM. Differences were determined using a Student’s t-test where p < 0.05 was significant (*); n = 6

### UT-A1/A3 KO mice have low urinary cAMP excretion despite elevated AVP levels

Serum analysis indicated that circulating AVP levels were significantly elevated in UT-A1/A3 KO mice (Fig. 4a). As previously discussed, AVP activates V2R, increasing intracellular cAMP levels (Hoffert et al., 2006; Pearce et al., 2014). With this mechanism in mind, we investigated urinary levels of cAMP. Interestingly, urinary cAMP levels were significantly lower in UT-A1/A3 KO mice as compared to WT animals, despite elevated AVP levels (Fig. 4b).

**Fig. 4.**
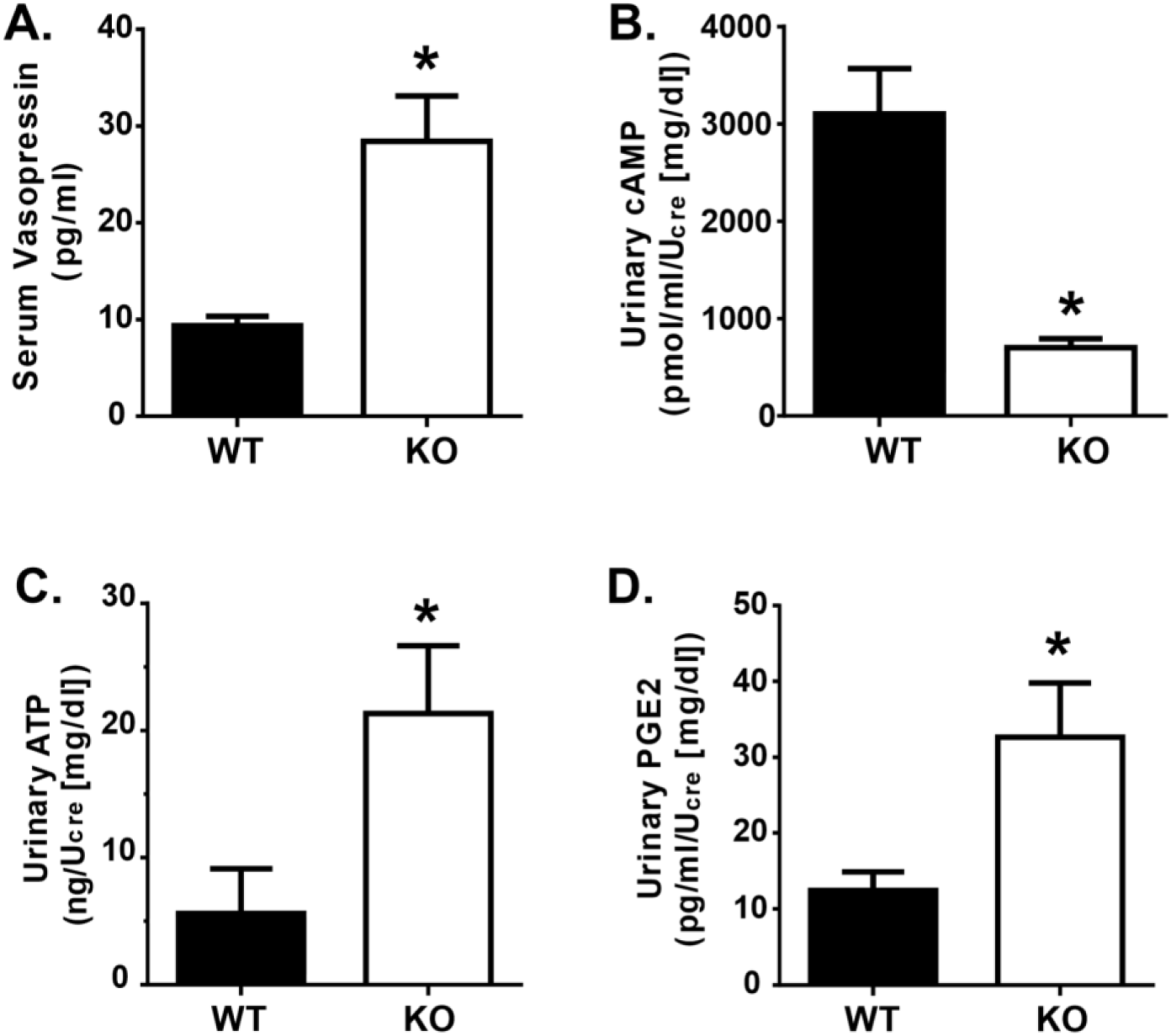
UT-A1/A3 KO mice excrete low amounts of cAMP, ATP, and PGE_2_ despite high circulating AVP. **a** Vasopressin was measured in serum samples collected from wild-type mice (WT) and UT-A1/A3 KO mice (KO). Mice were housed in metabolic cages and urine was collected over 24 hours. Fresh urine was then analyzed for **b** cAMP, **c** ATP, and **d** PGE_2_. All concentrations were normalized to creatinine (U_cre_) to correct for differences in urine dilution. All data presented as means ± SEM. Differences were determined using a Student’s t-test where p < 0.05 was significant (*); n = 4

AVP-mediated water reabsorption is inhibited by ATP-mediated activation of P2Y_2_, thereby dampening cAMP synthesis (Kishore et al., 1995). Given the decreased urinary cAMP levels observed in UT-A1/A3 KO mice, we suspected that there was an increase in ATP-mediated P2Y_2_ activation. Analysis confirms that urinary ATP levels were higher in UT-A1/A3 KO mice as compared to WT mice (Fig. 4c).

This elevation in urinary ATP led us to speculate that, similar to other animal models of AVP-resistant polyuria (Sun et al., 2005a), purinergic signaling was increased in the UT-A1/A3 KO mice. Although the IMCD releases moderate amounts of PGE_2_ under basal conditions, specific activation of the P2Y_2_ receptor results in an enhanced release of PGE_2_ from IMCD cells in a time- and concentration-dependent manner (Welch et al., 2003). Therefore, to indirectly analyze the ATP-activation of the P2Y_2_ receptor in the IMCD, we measured urinary excretion of PGE_2_. Immunoassays revealed that PGE_2_ levels were higher in UT-A1/A3 KO mice as compared to WT mice (Fig. 4d).

### UT-A1/A3 KO mice have elevated P2Y_2_ protein expression in the inner medulla

P2Y_2_ activation in response to ATP release in the collecting duct is critical for modulating water transport (Vallon et al., 2012). To determine if an increase in P2Y_2_ receptor abundance contributed to the observed increase in purinergic activity in UT-A1/A3 KO mice, we first measured P2Y_2_ mRNA expression in the medullary tissues of these animals using quantitative real-time RT-PCR (Fig 5). There was no significant difference in P2Y_2_ mRNA expression in the medullary tissues of WT and UT-A1/A3 KO mice (Fig 5).

**Fig. 5.**
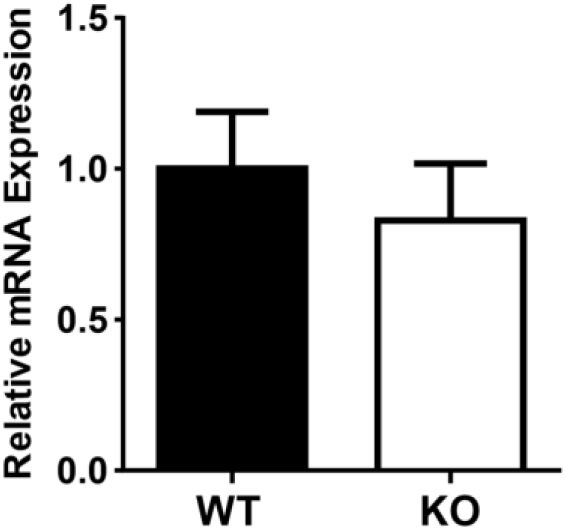
P2Y_2_ mRNA expression is unchanged in the medulla of UT-A1/A3 KO mice. Inner medullary tissues were dissected from control, wild-type mice (WT) and UT-A1/A3 KO (KO) to determine the expression of P2Y_2_ receptor mRNA by quantitative real-time RT-PCR. Results (means ± SEM) were computed as copies of P2Y_2_ receptor per copy of the housekeeping gene 18S (ΔΔCT_WT_, 3.73×10^−10^ ± 0.38; ΔΔCT_KO_, 0.30 ± 0.36). Differences in abundances were determined using a Student’s t-test where p < 0.05 was significant (*); n = 6

Others investigating the expression profiles of P2Y_2_ in rodent IMCD found that mRNA and protein levels of the receptor can be discordant (Kishore et al., 2005). Because of these observations, we measured P2Y_2_ protein abundance in the inner and outer medullary tissues of these animals.

Corroborating previous reports, we observed P2Y_2_ protein in both medullary tissues (Fig. 6) (Kishore et al., 2000). P2Y_2_ was more abundant in the inner medulla of UT-A1/A3 KO mice as compared to WT (Fig. 6), indicating that an increase in receptor abundance may be contributing to the observed increase in purinergic activity.

**Fig. 6.**
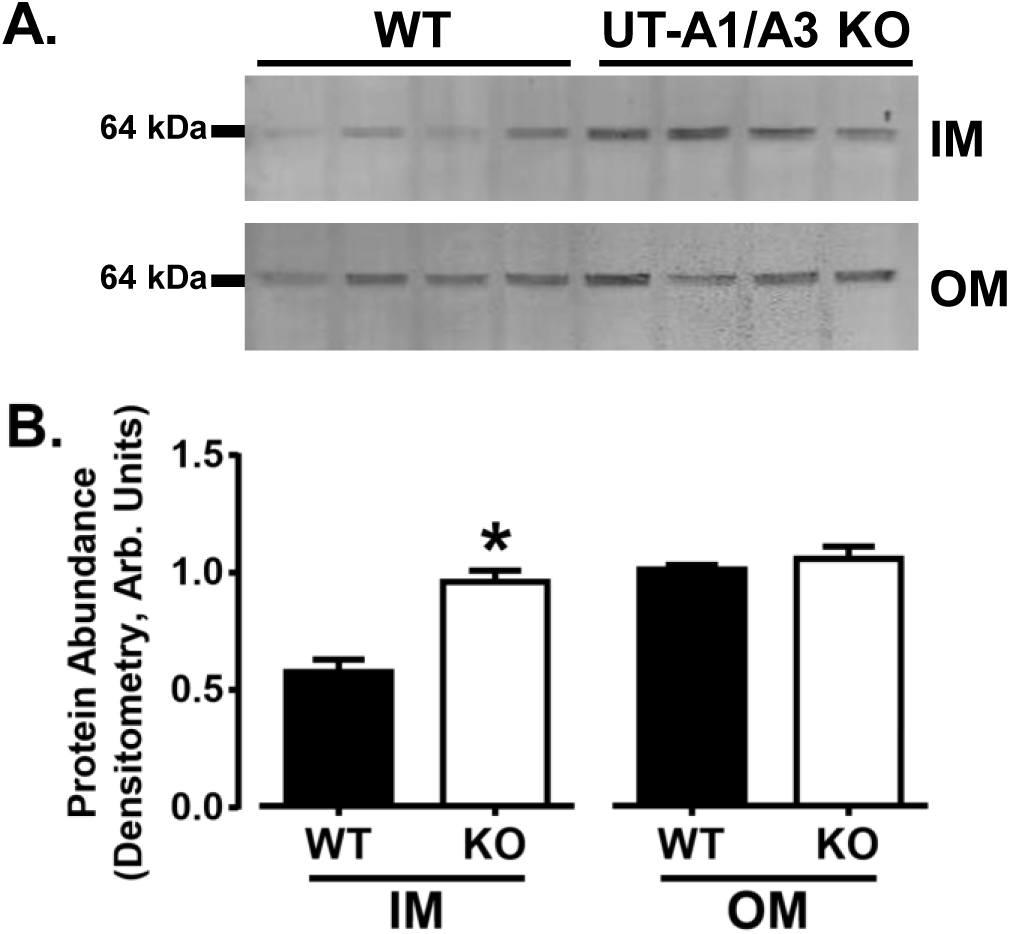
P2Y_2_ protein expression is elevated in the inner medulla of UT-A1/A3 KO mice. Inner medullary (IM) and outer medullary (OM) tissues were dissected from control, wild-type mice (WT) and UT-A1/A3 KO mice (KO), and subjected to western blot analysis. **a** Representative western blot images from tissues probed for P2Y_2_. Each lane represents one animal. **b** Normalized protein abundance quantification by band densitometry presented as means ± SEM. Differences in abundances were determined using a Student’s t-test where p < 0.05 was significant (*); n = 4

## Discussion

Many have shown that local factors, including ATP, may modulate the actions of AVP (Pearce et al., 2014). In the IMCD, ATP activates P2Y_2_ receptor, thereby inhibiting AVP-mediated water permeability via phospholipase C inhibition of cAMP. The activation of the P2Y_2_ receptor ultimately leads to PGE_2_ release (Sun et al., 2005a; Welch et al., 2003). ATP modulation of AVP-stimulated water transport in the IMCD has been studied extensively employing both pharmacological approaches (Ecelbarger et al., 1994; Kishore et al., 1995; Sun et al., 2005a; Sun et al., 2005b), and metabolic examination of mice lacking P2Y_2_ receptors (Rieg et al., 2007; Zhang et al., 2008); however, the role that urea transport plays in the purinergic-prostanoid system has not been extensively studied.

In the studies described herein, we found that mice simultaneously lacking both the apical urea transporter, UT-A1, and the basolateral urea transporter, UT-A3, have extensive polyuria despite unchanged levels of AQP2 expression and elevated circulating AVP. We observed lower levels of urinary cAMP in UT-A1/A3 KO mice as compared to wild-type litter mates. Furthermore, we observed that UT-A1/A3 KO mice have increased urinary PGE_2_ levels as compared to wild-type litter mates, as well as elevated expression of the P2Y_2_ receptor in inner medullary tissues. These data collectively indicate an increase in P2Y_2_ activity in UT-A1/A3 KO mice. Two aspects of IMCD physiology appear to be contributing to this observed change: 1) an increase in P2Y_2_ receptor protein abundance, and 2) the increase of the P2Y_2_ agonist ATP.

An increase in P2Y_2_ protein abundance, given suitable agonist levels, might lead to an increase in total P2Y_2_ activity; we observed, via western blot, an increase in total P2Y_2_ protein in the inner medulla. Previous studies have linked UT-A1 and P2Y_2_ abundance in a backwards fashion, demonstrating that in the absence of P2Y_2_, UT-A1 protein abundance is increased (Zhang et al., 2008). Additionally, in models of heightened purinergic activity, UT-A1 protein abundance is reduced (Blount et al., 2010; Zhang et al., 2009). Our data, in conjunction with previous data, indicates some sort of co-regulatory function between these proteins. Future studies are needed to clarify this relationship between the urea transporters and P2Y_2_ in the IMCD.

Although P2Y_2_ appears to regulate urea transporter abundance, it is unclear what mechanism is at play by which the absence of UT-A1 and UT-A3 stimulates P2Y_2_ protein synthesis. A likely explanation is that in the absence of urea transport in the IMCD, medullary tonicity is decreased, resulting in enhanced P2Y_2_ abundance. In vivo studies suggest that alterations in P2Y_2_ abundance are regulated by medullary tonicity (Kishore et al., 2005), which was later supported by in vitro studies in mpkCCD cells (Wildman et al., 2009). In fact, inner medullary tissue osmolality is significantly higher in mice lacking P2Y_2_ receptors, suggesting that the elevated UT-A1 abundance observed in these same mice increases their urea recycling capability (Zhang et al., 2013; Zhang et al., 2008).

Interestingly, we observed a discrepancy between P2Y_2_ protein abundance and mRNA expression, but discrepancies of this sort have been previously reported (Insel et al., 2001; Kishore et al., 2005). It is not uncommon for G protein-coupled receptors like P2Y_2_ to be sequestered into the cytoplasm to either be recycled back to the plasma membrane, or sorted for degradation into lysosomes. Biochemical studies examining the cellular trafficking of a tagged-P2Y_2_ demonstrate that the receptor can be internalized by a clathrin-mediated pathway and degraded through a proteasome pathway (Sromek and Harden, 1998; Tulapurkar et al., 2005). Increased AVP levels induce a cAMP-dependent translocation of P2Y_2_ receptors into the basolateral membrane (Wildman et al., 2009) suggesting that the observed receptor recycling and possibly degradation may be mediated by AVP. This and other possible explanations will need to be explored in more extensive investigations to elucidate the reason for the observed difference; other explanations may involve posttranslational modification, or other aspects of P2Y_2_ protein stability and turnover.

Increased levels of ATP might also drive increased P2Y_2_ activation, and we observed an increase of urinary ATP in UT-A1/A3 KO mice. As previously discussed, rapid changes to the tonicity of the inner medullary interstitium is responsible for increased IMCD cell size, and the subsequent release of ATP is essential to normalizing cell volume by driving P2Y_2_ activation. Previous studies indicate that acute water loading increases the release of ATP due to the reduction of extracellular tonicity and a resultant increase in cell volume (Rieg et al., 2007). It has been previously demonstrated, and we have confirmed, that UT-A1/A3 KO mice have a reduced ability to reabsorb urea, and thus have higher levels of urinary urea when compared to wild type animals. Given that urea is a major osmotic player, its presence in luminal fluid could be building osmotic potential toward the lumen of the collecting duct. Water moving from the interstitium toward the collecting duct, through the basolateral aquaporins AQP3 and AQP4, could increase cell volume, releasing ATP at both the basolateral and apical membranes, contributing to the observed increase in ATP, and ultimately to an increase in P2Y_2_ activation.

Among previously mentioned stimulators, increased luminal flow increases luminal ATP expression in principal cells of the collecting duct (Jensen et al., 2007; Praetorius and Leipziger, 2010). UT-A1/A3 KO mice are polyuric, and as such, luminal flow rates in the collecting duct are likely increased as compared to wild-type mice. It is worth noting, however, that urinary flow rate alone is not a good predictor of urinary ATP excretion (Rieg and Vallon, 2009). At this time, it is unclear what other factors may be involved.

We observed that AQP2 protein abundance remains unchanged in UT-A1/A3 KO mice, and as such the potential for water movement is likely unaffected. Granted that urea recycling in the inner medulla is ablated in the UT-A1/A3 KO mice, there are no reported differences in the expression of sodium transporters in the distal nephron and collecting ducts of these animals (Fenton et al., 2005), suggesting that sodium reabsorption can still contribute to generating a weakened osmotic gradient in the medullary interstitium. However, because of possible elevated purinergic-prostanoid activity in the UT-A1/A3 KO mice, we predict that any possible water reabsorption by AQP2 would be inhibited despite the presence of a dampened osmotic gradient. Therefore, the polyuria seen in the UT-A1/A3 KO mice is likely compounded by the inability of these mice to reabsorb luminal water in the IMCD.

While many studies agree that PGE_2_ plays an important role in regulating water excretion by modulating the effects of AVP, PGE_2_-diuretic effects are extremely variable. For instance, PGE_2_ stimulates water transport in the absence of AVP, but decreases water and urea permeability in the presence of AVP (Hebert et al., 1993; Rouch and Kudo, 2000). Although the understanding of the underlying molecular mechanisms behind the PGE_2_-mediated inhibition of urea and water transport remains incomplete, the differences in PGE2-mediated actions are likely explained by the diverse range of biologic actions elicited by the four E-prostanoid receptor subtypes, EP1, EP2, EP3, and EP4 (Breyer and Breyer, 2001; Olesen and Fenton, 2013). While not investigated herein, we speculate that the elevated PGE_2_ levels observed in UT-A1/A3 KO mice may inhibit AVP-induced water reabsorption through the EP3 receptor. EP3 receptor stimulation, which is coupled to G_*i*_, would explain the decrease levels of urinary cAMP in these mice despite elevated AVP. EP3-deficient mice can concentrate urine similar to WT mice after both water loading and water deprivation; however, pharmacological inhibition of PGE_2_ production only increases urine osmolality in WT mice. These studies suggest that PGE_2_ action through the EP3 receptor is not essential for urine concentration under normal conditions, but when PGE_2_ is elevated, as observed in the UT-A1/A3 KO mice, the EP3 receptor likely plays a crucial role (Fleming et al., 1998). Further support comes from a recent study demonstrating that lithium-induced polyuria is attenuated in P2Y_2_ KO mice (Zhang et al., 2013). EP3 receptor expression is downregulated in the collecting ducts of lithium-treated P2Y_2_ KO mice, and PGE_2_-stimulation of isolated IMCD tubules from these mice stimulates cAMP production at higher rates than lithium-treated WT mice. Not only do these findings suggest that the IMCD can become more sensitive to PGE_2_ in response to altered physiological settings like elevated urine flow, but these studies also demonstrate that P2Y_2_ is a prominent regulator of EP3 abundance.

In conclusion, using UT-A1/A3 KO mice we have demonstrated that urea transport in the inner medulla is modulated in some part by the purinergic-prostanoid system. Although most previous studies have focused on local regulation of water reabsorption in the IMCD by ATP, the effect of this nucleotide on urea handling in the inner medulla is largely unknown. This work represents the first step in understanding the connection between P2Y_2_-regulation of the urea transporters UT-A1 and UT-A3.

## Acknowledgements

The authors would like to thank Jessie Kuo for technical assistance, and Dr. Jeff M. Sands for generously providing UT-A1/A3 KO mice.

## References

Blount, M. A., Klein, J. D., Martin, C. F., Tchapyjnikov, D. and Sands, J. M. (2007). Forskolin stimulates phosphorylation and membrane accumulation of UT-A3. Am J Physiol Renal Physiol 293, F1308–F1313.

Blount, M. A., Mistry, A. C., Fröhlich, O., Price, S. R., Chen, G., Sands, J. M. and Klein, J. D. (2008). Phosphorylation of UT-A1 urea transporter at serines 486 and 499 is important for vasopressin-regulated activity and membrane accumulation. Am J Physiol Renal Physiol 295, F295–F299.

Blount, M. A., Sim, J. H., Zhou, R., Martin, C. F., Lu, W., Sands, J. M. and Klein, J. D. (2010). Expression of transporters involved in urine concentration recovers differently after cessation of lithium treatment. Am J Physiol Renal Physiol 298, F601–8.

Boone, M. and Deen, P. M. T. (2008). Physiology and pathophysiology of the vasopressin-regulated renal water reabsorption. Pflugers Archiv 456, 1005–1024.

Breyer, M. D. and Breyer, R. M. (2001). G protein-coupled prostanoid receptors and the kidney. Annu Rev Physiol 63, 579–605.

Burnstock, G., Evans, L. C. and Bailey, M. A. (2014). Purinergic signalling in the kidney in health and disease. Purinergic Signal 10, 71–101.

Chabardes-Garonne, D., Mejean, A., Aude, J. C., Cheval, L., Di Stefano, A., Gaillard, M. C., Imbert-Teboul, M., Wittner, M., Balian, C., Anthouard, V. et al. (2003). A panoramic view of gene expression in the human kidney. Proc Natl Acad Sci U S A 100, 13710–5.

Chou, C.-L., Yu, M.-J., Kassai, E. M., Morris, R. G., Hoffert, J. D., Wall, S. M. and Knepper, M. A. (2008). Roles of basolateral solute uptake via NKCC1 and of myosin II in vasopressin-induced cell swelling in inner medullary collecting duct. American Journal of Physiology. Renal Physiology 295, F192–F201.

Ecelbarger, C. A., Maeda, Y., Gibson, C. C. and Knepper, M. A. (1994). Extracellular ATP increases intracellular calcium in rat terminal collecting duct via a nucleotide receptor. Am J Physiol Renal Physiol 267, F998–F1006.

Edwards, R. M. (2002). Basolateral, but not apical, ATP inhibits vasopressin action in rat inner medullary collecting duct. European Journal of Pharmacology 438, 179–181.

Erb, L., Liao, Z., Seye, C. and Weisman, G. (2006). P2 receptors: intracellular signaling. Pflugers Archiv 452, 552–562.

Fenton, R. A., Chou, C.-L., Stewart, G. S., Smith, C. P. and Knepper, M. A. (2004). Urinary concentrating defect in mice with selective deletion of phloretin-sensitive urea transporters in the renal collecting duct. Proc Natl Acad Sci U S A 101, 7469–7474.

Fenton, R. A., Flynn, A., Shodeinde, A., Smith, C. P., Schnermann, J. and Knepper, M. A. (2005). Renal phenotype of UT-A urea transporter knockout mice. J Am Soc Nephrol 16, 1583–92.

Fleming, E. F., Athirakul, K., Oliverio, M. I., Key, M., Goulet, J., Koller, B. H. and Coffman, T. M. (1998). Urinary concentrating function in mice lacking EP3 receptors for prostaglandin E2. Am J Physiol 275, F955–61.

Ganote, C. E., Grantham, J. J., Moses, H. L., Burg, M. B. and Orloff, J. (1968). Ultrastructural studies of vasopressin effect on isolated perfused renal collecting tubules of the rabbit. J Cell Biol 36, 355–67.

Hebert, R. L., Jacobson, H. R., Fredin, D. and Breyer, M. D. (1993). Evidence that separate PGE2 receptors modulate water and sodium transport in rabbit cortical collecting duct. Am J Physiol 265, F643–50.

Hoffert, J. D., Pisitkun, T., Wang, G., Shen, R.-F. and Knepper, M. A. (2006). Quantitative phosphoproteomics of vasopressin-sensitive renal cells: Regulation of aquaporin-2 phosphorylation at two sites. Proceedings of the National Academy of Sciences 103, 7159–7164.

Hovater, M., Olteanu, D., Hanson, E., Cheng, N.-L., Siroky, B., Fintha, A., Komlosi, P., Liu, W., Satlin, L., Bell, P. D. et al. (2008). Loss of apical monocilia on collecting duct principal cells impairs ATP secretion across the apical cell surface and ATP-dependent and flow-induced calcium signals. Purinergic Signal 4, 155–170.

Ilori, T. O., Blount, M. A., Martin, C. F., Sands, J. M. and Klein, J. D. (2013). Urine concentration in the diabetic mouse requires both urea and water transporters. Am J Physiol Renal Physiol 304, F103–11.

Insel, P. A., Ostrom, R. S., Zambon, A. C., Hughes, R. J., Balboa, M. A., Shehnaz, D., Gregorian, C., Torres, B., Firestein, B. L., Xing, M. et al. (2001). Extracellular ATP and cAMP as Paracrine and Interorgan Regulators of Renal Function P2Y Receptors of MDCK Cells: Epithelial Cell Regulation by Extracellular Nucleotides. Clinical and Experimental Pharmacology and Physiology 28, 351–354.

Jankowski, M., Szamocka, E., Kowalski, R., Angielski, S. and Szczepanska-Konkel, M. (2011). The effects of P2X receptor agonists on renal sodium and water excretion in anaesthetized rats. Acta Physiol (Oxf) 202, 193–201.

Jensen, M. E. J., Odgaard, E., Christensen, M. H., Praetorius, H. A. and Leipziger, J. (2007). Flow-Induced [Ca2+]i Increase Depends on Nucleotide Release and Subsequent Purinergic Signaling in the Intact Nephron. Journal of the American Society of Nephrology 18, 2062–2070.

Kishore, B. K., Chou, C. L. and Knepper, M. A. (1995). Extracellular nucleotide receptor inhibits AVP-stimulated water permeability in inner medullary collecting duct. Am J Physiol 269, F863–9.

Kishore, B. K., Ginns, S. M., Krane, C. M., Nielsen, S. and Knepper, M. A. (2000). Cellular localization of P2Y(2) purinoceptor in rat renal inner medulla and lung. Am J Physiol Renal Physiol 278, F43–51.

Kishore, B. K., Krane, C. M., Miller, R. L., Shi, H., Zhang, P., Hemmert, A., Sun, R. and Nelson, R. D. (2005). P2Y2 receptor mRNA and protein expression is altered in inner medullas of hydrated and dehydrated rats: relevance to AVP-independent regulation of IMCD function. Am J Physiol Renal Physiol 288, F1164–72.

Klein, J. D., Blount, M. A., Fröhlich, O., Denson, C. E., Tan, X., Sim, J. H., Martin, C. F. and Sands, J. M. (2010). Phosphorylation of UT-A1 on serine 486 correlates with membrane accumulation and urea transport activity in both rat IMCDs and cultured cells. Am J Physiol Renal Physiol 298, F935–F940.

Klein, J. D., Blount, M. A. and Sands, J. M. (2011). Urea Transport in the Kidney. In Comprehensive Physiology: John Wiley & Sons, Inc.

Naruse, M., Klein, J. D., Ashkar, Z. M., Jacobs, J. D. and Sands, J. M. (1997). Glucocorticoids downregulate the vasopressin-regulated urea transporter in rat terminal inner medullary collecting ducts. Journal of the American Society of Nephrology 8, 517–23.

Olesen, E. T. B. and Fenton, R. A. (2013). Is There a Role for PGE2 in Urinary Concentration? Journal of the American Society of Nephrology 24, 169–178.

Pannabecker, T. L., Dantzler, W. H., Layton, H. E. and Layton, A. T. (2008). Role of three-dimensional architecture in the urine concentrating mechanism of the rat renal inner medulla. Am J Physiol Renal Physiol 295, F1271–F1285.

Pearce, D., Soundararajan, R., Trimpert, C., Kashlan, O. B., Deen, P. M. and Kohan, D. E. (2014). Collecting Duct Principal Cell Transport Processes and Their Regulation. Clin J Am Soc Nephrol.

Praetorius, H. A. and Leipziger, J. (2010). Intrarenal Purinergic Signaling in the Control of Renal Tubular Transport. Annual Review of Physiology 72, 377–393.

Rieg, T., Bundey, R. A., Chen, Y., Deschenes, G., Junger, W., Insel, P. A. and Vallon, V. (2007). Mice lacking P2Y2 receptors have salt-resistant hypertension and facilitated renal Na+ and water reabsorption. Faseb j 21, 3717–26.

Rieg, T., Tang, T., Murray, F., Schroth, J., Insel, P. A., Fenton, R. A., Hammond, H. K. and Vallon, V. (2010). Adenylate Cyclase 6 Determines cAMP Formation and Aquaporin-2 Phosphorylation and Trafficking in Inner Medulla. Journal of the American Society of Nephrology 21, 2059–2068.

Rieg, T. and Vallon, V. (2009). ATP and adenosine in the local regulation of water transport and homeostasis by the kidney. Am J Physiol Regul Integr Comp Physiol 296, R419–27.

Rouch, A. J. and Kudo, L. H. (2000). Role of PGE(2) in alpha(2)-induced inhibition of AVP-and cAMP-stimulated H(2)O, Na(+), and urea transport in rat IMCD. Am J Physiol Renal Physiol 279, F294–301.

Rouse, D., Leite, M. and Suki, W. N. (1994). ATP inhibits the hydrosmotic effect of AVP in rabbit CCT: evidence for a nucleotide P2u receptor. Am J Physiol Renal Physiol 267, F289–F295.

Sands, J. M. (2003). Molecular Mechanisms of Urea Transport. The Journal of Membrane Biology 191, 149–163.

Sands, J. M., Blount, M. A. and Klein, J. D. (2011). Regulation of renal urea transport by vasopressin. Trans Am Clin Climatol Assoc 122, 82–92.

Sands, J. M. and Layton, H. E. (2009). The Physiology of Urinary Concentration: An Update. Seminars in Nephrology 29, 178–195.

Schwiebert, E. M. (2001). ATP release mechanisms, ATP receptors and purinergic signalling along the nephron. Clin Exp Pharmacol Physiol 28, 340–50.

Schwiebert, E. M. and Kishore, B. K. (2001). Extracellular nucleotide signaling along the renal epithelium. Am J Physiol Renal Physiol 280, F945–F963.

Sim, J. H., Himmel, N. J., Redd, S. K., Pulous, F. E., Rogers, R. T., Black, L. N., Hong, S. M., von Bergen, T. N. and Blount, M. A. (2014). Absence of PKC-alpha attenuates lithium-induced nephrogenic diabetes insipidus. PLoS One 9, e101753.

Sromek, S. M. and Harden, T. K. (1998). Agonist-induced internalization of the P2Y2 receptor. Mol Pharmacol 54, 485–94.

Sun, R., Carlson, N. G., Hemmert, A. C. and Kishore, B. K. (2005a). P2Y2 receptor-mediated release of prostaglandin E2 by IMCD is altered in hydrated and dehydrated rats: relevance to AVP-independent regulation of IMCD function. Am J Physiol Renal Physiol 289, F585–92.

Sun, R., Miller, R. L., Hemmert, A. C., Zhang, P., Shi, H., Nelson, R. D. and Kishore, B. K. (2005b). Chronic dDAVP infusion in rats decreases the expression of P2Y2 receptor in inner medulla and P2Y2 receptor-mediated PGE2 release by IMCD. Am J Physiol Renal Physiol 289, F768–76.

Tulapurkar, M. E., Schafer, R., Hanck, T., Flores, R. V., Weisman, G. A., Gonzalez, F. A. and Reiser, G. (2005). Endocytosis mechanism of P2Y2 nucleotide receptor tagged with green fluorescent protein: clathrin and actin cytoskeleton dependence. Cell Mol Life Sci 62, 1388–99.

Turner, C. M., Vonend, O., Chan, C., Burnstock, G. and Unwin, R. J. (2003). The pattern of distribution of selected ATP-sensitive P2 receptor subtypes in normal rat kidney: an immunohistological study. Cells Tissues Organs 175, 105–17.

Unwin, R. J., Bailey, M. A. and Burnstock, G. (2003). Purinergic signaling along the renal tubule: the current state of play. News Physiol Sci 18, 237–41.

Vallon, V. and Rieg, T. (2011). Regulation of renal NaCl and water transport by the ATP/UTP/P2Y2 receptor system. Am J Physiol Renal Physiol 301, F463–75.

Vallon, V., Stockand, J. and Rieg, T. (2012). P2Y receptors and kidney function. Wiley Interdiscip Rev Membr Transp Signal 1, 731–742.

Welch, B. D., Carlson, N. G., Shi, H., Myatt, L. and Kishore, B. K. (2003). P2Y2 receptor-stimulated release of prostaglandin E2 by rat inner medullary collecting duct preparations. Am J Physiol Renal Physiol 285, F711–21.

Wildman, S. S., Boone, M., Peppiatt-Wildman, C. M., Contreras-Sanz, A., King, B. F., Shirley, D. G., Deen, P. M. and Unwin, R. J. (2009). Nucleotides downregulate aquaporin 2 via activation of apical P2 receptors. J Am Soc Nephrol 20, 1480–90.

Zhang, Y., Li, L., Kohan, D. E., Ecelbarger, C. M. and Kishore, B. K. (2013). Attenuation of lithium-induced natriuresis and kaliuresis in P2Y2 receptor knockout mice. Am J Physiol Renal Physiol 305, F407–16.

Zhang, Y., Nelson, R. D., Carlson, N. G., Kamerath, C. D., Kohan, D. E. and Kishore, B. K. (2009). Potential role of purinergic signaling in lithium-induced nephrogenic diabetes insipidus. Am J Physiol Renal Physiol 296, F1194–201.

Zhang, Y., Sands, J. M., Kohan, D. E., Nelson, R. D., Martin, C. F., Carlson, N. G., Kamerath, C. D., Ge, Y., Klein, J. D. and Kishore, B. K. (2008). Potential role of purinergic signaling in urinary concentration in inner medulla: insights from P2Y2 receptor gene knockout mice. Am J Physiol Renal Physiol 295, F1715–24.

